# metGWAS 1.0: An R workflow for network-driven over-representation analysis between independent metabolomic and meta-genome wide association studies

**DOI:** 10.1101/2022.08.09.503325

**Authors:** Saifur R. Khan, Andreea Obersterescu, Erica P. Gunderson, Michael B. Wheeler, Brian J. Cox

**Affiliations:** Department of Physiology, University of Toronto, Ontario, Canada; Advanced Diagnostics, Metabolism, Toronto General Research Institute, Ontario, Canada; University of Pittsburgh Medical Center, Pittsburgh, Pennsylvania, USA; Kaiser Permanente Northern California, Division of Research, Oakland, CA, USA; Kaiser Permanente Bernard J. Tyson School of Medicine, Pasadena, CA, USA; Department of Obstetrics and Gynaecology, University of Toronto, Ontario, Canada

**Keywords:** Genome-wide association studies (GWAS), metabolomics-GWAS, standalone metabolomics, genetic predisposition, risk-alleles, gene loci, GWAS-network model, Jaccard coefficient

## Abstract

**Background:** Many diseases may result from disrupted metabolic regulation. Metabolite-GWAS studies assess the association of polymorphic variants with metabolite levels in body fluids. While these studies are successful, they have a high cost and technical expertise burden due to combining the analytical biochemistry of metabolomics with the computational genetics of GWAS. Currently, there are 100s of standalone metabolomics and GWAS studies related to similar diseases or phenotypes. A method that could statically evaluate these independent studies to find novel metabolites-genes association is of high interest. Although such an analysis is limited to genes with known metabolite interactions due to the unpaired nature of the data sets, any discovered associations may represent biomarkers and druggable targets for treatment and prevention.

**Methods:** We developed a bioinformatics tool, metGWAS 1.0, that generates and statistically compares metabolic and genomic gene sets using a hypergeometric test. Metabolic gene sets are generated by mapping disease-associated metabolites to interacting proteins (genes) via online databases. Genomic gene sets are identified from a network representation of the GWAS Catalog comprising 100s of studies.

**Results:** The metGWAS 1.0 tool was evaluated using standalone metabolomics datasets extracted from two metabolomics-GWAS case studies. In case-study 1, a cardiovascular disease association study, we identified nine genes (APOA5, PLA2G5, PLA2G2D, PLA2G2E, PLA2G2F, LRAT, PLA2G2A, PLB1, and PLA2G7) that interact with metabolites in the KEGG glycerophospholipid metabolism pathway and contain polymorphic variants associated with cardiovascular disease (*P* < 0.005). The gene APOA5 was matched from the original metabolomics-GWAS study. In case study 2, a urine metabolome study of kidney metabolism in healthy subjects, we found marginal significance (*P = 0*.*10 and P = 0*.*13*) for glycine, serine, and threonine metabolism and alanine, aspartate, and glutamate metabolism pathways to GWAS data relating to kidney disease.

**Conclusion:** The metGWAS 1.0 platform provides insight into developing methods that bridge standalone metabolomics and disease and phenotype GWAS data. We show the potential to reproduce findings of paired metabolomics-GWAS data and provide novel associations of gene variation and metabolite expression.

## INTRODUCTION

Biochemical reactions catalyzed by enzymes produce different small molecules, named metabolites[1, 2], representing the complex integration of biological states with environmental and lifestyle factors[3]. These metabolites interact with diverse proteins and serve critical roles as nutrients, building blocks, receptor ligands, transcriptional cofactors, genetic activators, and suppressors that modulate the biological systems’ adaptive responses [4]. In many diseases, there is a disruption of metabolic homeostasis. In particular, type 2 diabetes and cardiovascular disease often show metabolic changes before their onset [3, 5-10]. Recent advances in high throughput methods to profile bodily fluids and tissue metabolomes have provided a wealth of information regarding metabolic health. Robust methods, including mass spectrometry and nuclear magnetic resonance, that quantify the metabolome are expanding and becoming universal tools to identify disease pathology [11]. This is greatly facilitated by the ample availability of blood plasma, saliva, and urine which are easily obtained and contain a rich diversity of metabolites[12].

Genome-wide association studies (GWAS) are critical in discovering genetic predispositions (i.e., disease-risk loci and risk-alleles) for most diseases [13]. However, many disease variants have small effects and are difficult to detect, even with large cohorts of individuals. On the other hand, variant associations with metabolite levels are stronger [14]. The integrated method, metabolomics-GWAS, aims to discover the genetic predispositions for metabolic alterations that may lead to disease [15, 16]. While a successful strategy, the combined technological complexity and high cost are barriers to expanded use of metabolomics-GWAS. Notably, many standalone metabolomics and GWAS studies investigating the same diseases exist in independent cohorts. An *in silico* platform that allows the merging of independent GWAS and metabolomics data to identify statistical relationships could facilitate the discovery of new target candidates for diagnostic and therapeutic interventions.

Motivated to fill this need, we created metGWAS 1.0, a standalone R pipeline that integrates independent GWAS and metabolomics data sets using a network-based systems biology approach. Enriched metabolites are mapped to catalyzing and interacting protein-coding genes through the Human Metabolome Database ((https://hmdb.ca/) and the UniProt database (www.uniprot.org). A network representation of the GWAS Catalog, which consolidates 100s of published GWAS studies, is used to identify metabolite interacting genes possibly related to the phenotype of interest. Lastly, a hypergeometric test tests the observed intersection of metabolite and genomic data sets.

## IMPLEMENTATION

This workflow utilizes MetaboAnalyst tools and integration into custom functions within *R* provided as metGWAS 1.0 (**Figure-1**). The user does not need any prior knowledge of *R*. We include the R-script for running the workflow and a user manual with a tutorial and technical information (**supplementary material 1**). To facilitate mapping metabolites to genes, we limit data to only those genes coding for proteins annotated to interact with a metabolite through the Human Metabolome Database ((https://hmdb.ca/). This workflow has been described below step-by-step.

**Figure 1:**
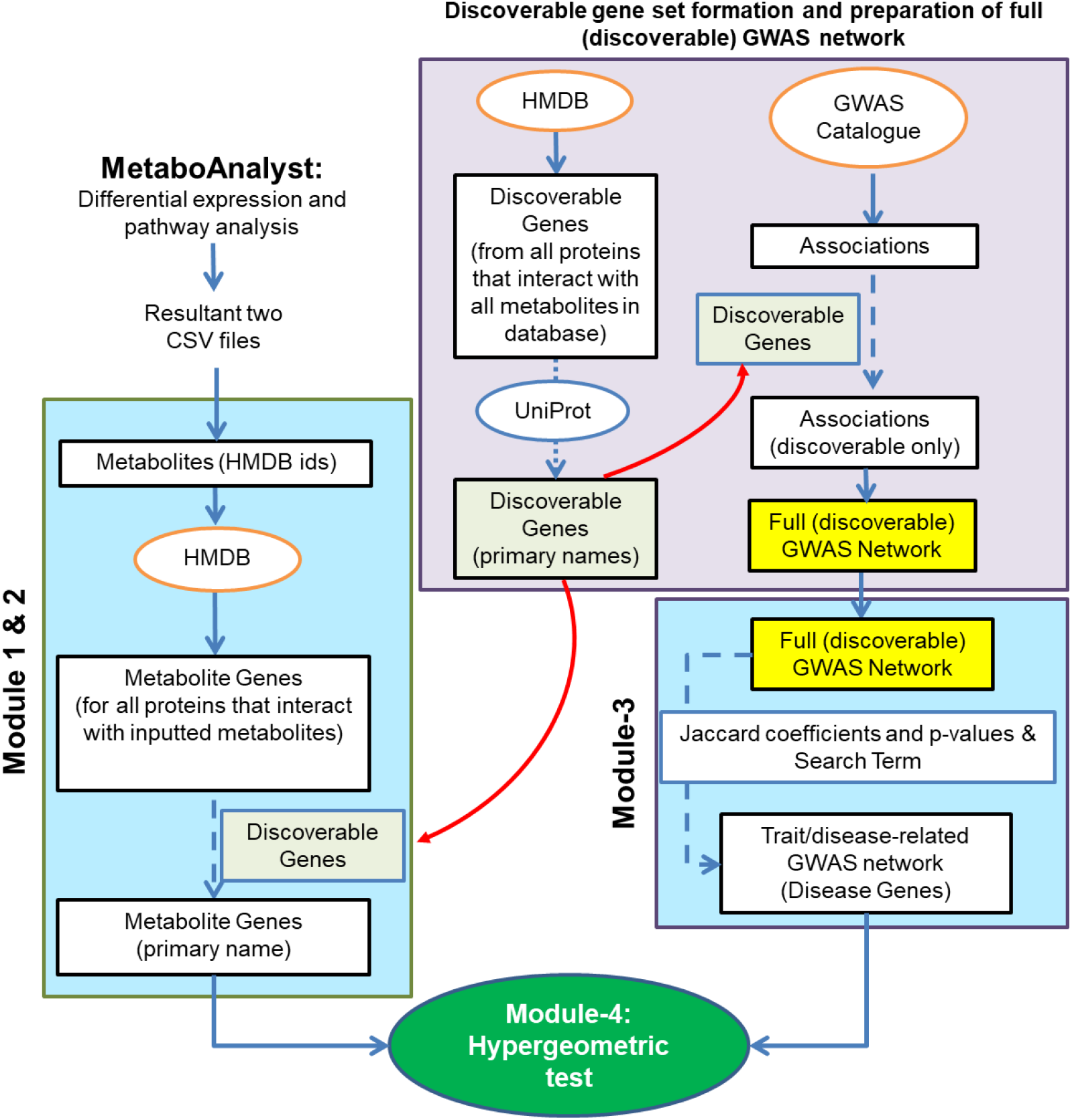
A diagram of the workflow for metGWAS 1.0 analysis. MetaboAnalyst calculates enriched metabolites and KEGG pathways. Modules 1 and 2 convert enriched KEGG pathways into interacting proteins and genes. Module 3 filters the network model of the GWAS Catalog. Module 4 performs a hypergeometric test comparing the enriched metabolite interacting genes with the trait/disease-related GWAS network. Saved objects provided with the workflow are highlighted yellow and the steps taken to create them are in the pink boxes. The steps the workflow runs every time are indicated by blue boxes.

### MetaboAnalyst: Differential expression and pathway analysis

The first step of the workflow starts with differential expression analysis of an appropriately preprocessed (i.e., missing value estimation, normalization, and data transformation) standalone metabolomic dataset through the MetaboAnalyst platform (https://www.metaboanalyst.ca/), a well-known bioinformatics platform for metabolomics analysis. The differentially expressed metabolites are mapped to human metabolite database (HMDB) identifiers and linked to the *Homo sapiens* KEGG pathways library of MetaboAnalyst. Next, hypergeometric and relative-betweenness centrality tests identify any over-represented KEGG pathways. MetaboAnalyst will generate two output CSV files, named ‘name_map’ and ‘pathway_results’, which will be required when initiating module-1 of this metGWAS 1.0 platform.

We recommend separating downregulated and upregulated metabolites’ HMBD IDs to identify downregulated and upregulated KEGG pathways.

### Module-1: Identification and annotation of the corresponding metabolites for the selected over-represented KEGG pathways

In module-1, using the CSV files ‘name_map’ and ‘pathway_results’, the user specifies an FDR p-value threshold and impact (the relative-betweenness centrality, valued from 0 to 1) to filter the list of MetaboAnalyst identified KEGG pathways. We recommend using FDR ≤ 0.05 to select the significantly altered KEGG pathways. Once the KEGG pathway(s) of interest is selected, module-1 annotates metabolites belonging to the chosen KEGG pathway using their HMDB ids through a call-out to the KEGG database. Large numbers of pathways (12s) and metabolites (100s) can take an extended processing time (20-30 minutes).

### Module-2: Identification of the metabolite-interacting proteins and their gene symbols

Module-2 maps metabolites to interacting proteins by searching metabolites’ HMDB ids in the HMDB database (https://hmdb.ca/). The list of interacting proteins is annotated as UniProt IDs (https://www.uniprot.org/) and mapped to current gene symbols. Only human genes and associated gene symbols are used as the GWAS Catalog is based on human studies.

### Module-3: trait/disease-related GWAS network preparation from the GWAS Catalog

The entire GWAS Catalog, a GWAS database, is filtered to only include genes with annotated metabolite interactions through the Human Metabolome Database ((https://hmdb.ca/). We provide this filtered GWAS Catalog network as a prebuilt network (**supplementary material 1)**. We provide code to rebuild the network using updated GWAS catalogues, which takes 3-4 hours on a standard computer. If the user chooses to update the GWAS network, they must download an updated version of the GWAS Catalog, which the workflow will use to create a network.

To construct the GWAS network (as of December 2019), each study/trait from the table is used as a node in the network. The nodes are annotated with PubMed ID, study title, study trait (phenotype) and -the reported significant genes. Next, the genes annotated to the nodes are filtered to contain only those found in the HMDB (referred to as the discoverable gene set) (**Figure-1**, detail is given in the technical document in **supplementary material-1**). Filtering to include only metabolite interacting genes ensures that both gene sets (metabolic and GWAS) are derived from the same background set of genes. To build edges between nodes, the Jaccard coefficient (a scoring system that measures similarity between two sets of objects) and associated p-values are calculated between all pairwise nodes. Significant edges (based on user-supplied thresholds for Jaccard coefficients and p-values) are kept.

A keyword search of the GWAS catalogue identifies primary nodes annotated with the keyword as part of the study title or the reported trait. This process is illustrated in **Figures 2 & 3**. Next, the 1^st^-degree neighbour nodes are selected to ensure GWAS studies with related phenotypes are captured. The workflow allows the user to set thresholds for Jaccard coefficients and their FDR-adjusted p-values for nearest neighbour selection. We considered overlap significant if the Jaccard coefficient ≥ 0.5 (a Jaccard coefficient of 0.5 is considered very conservative) and the FDR-corrected *p*-value ≤ 0.05. A lower Jaccard coefficient value (<0.5) increases the chance of network inflation with unrelated genes and studies, increasing the false discovery. The details of the construction of the trait/disease-related GWAS network (or disease gene set/network) are illustrated in **Figure 3**.

**Figure 2:**
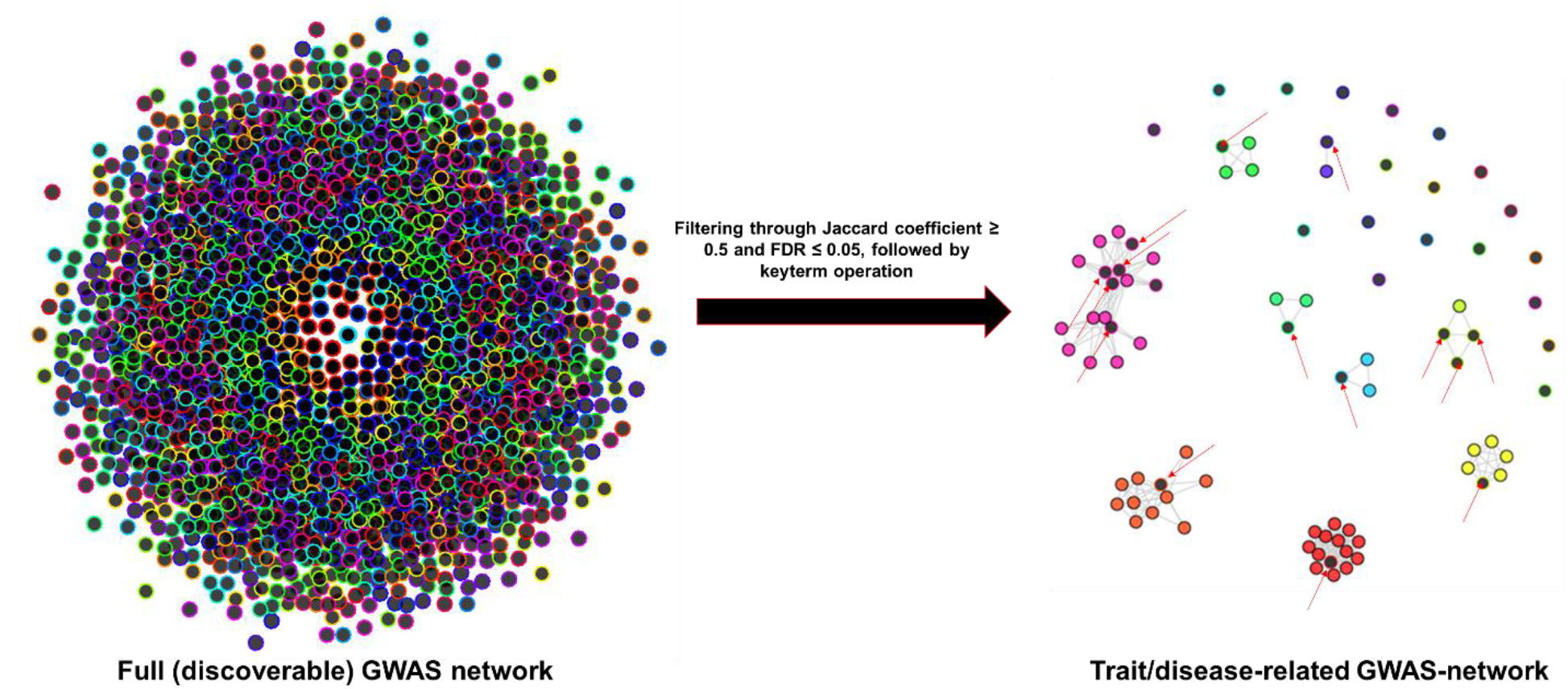
A representation of a trait/disease-related GWAS network created by module-3 from the full (discoverable) GWAS network. Each node represents a study/trait. Nodes are linked when there is an overlap between their associated genes. After filtering, black denotes a primary node found by the key term search (highlighted here with red arrows). Nearest neighbors are coloured instead of black. Here, this representative trait/disease-related GWAS network was constructed using case study-2.

**Figure 3:**
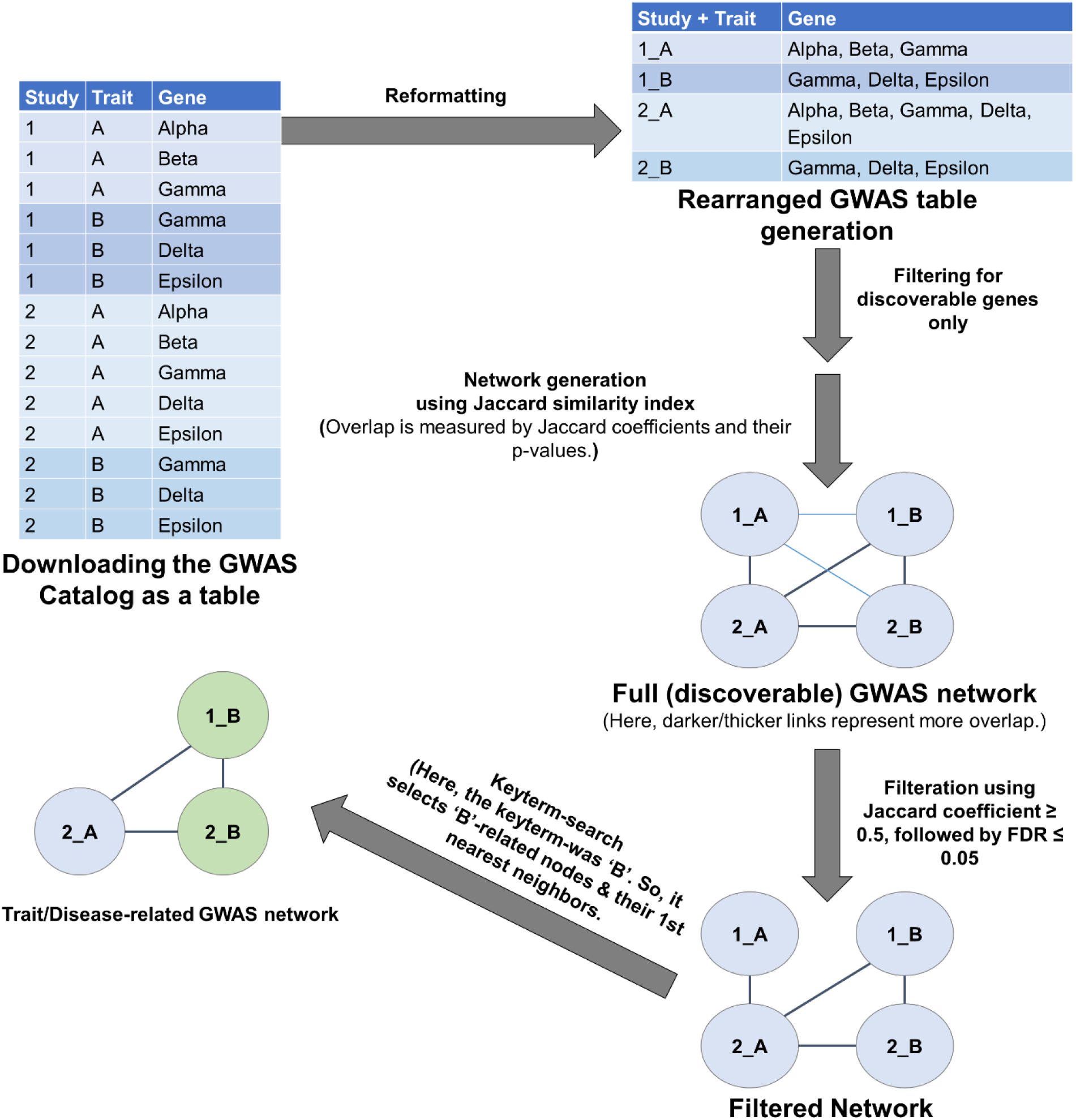
Schema of trait/disease-related GWAS-network creation. The GWAS Catalog is downloaded as a table (represented by the table on the top left). The table includes information on PubMed ids, study titles, reported traits, SNPs, and mapped genes. We work with the mapped genes, not directly with the SNPs. The table is then collapsed into a smaller table (top right). Here, each study+trait combination seen in the top-left table represents a unique row in the new table where all the gene associations have been listed together. Each row in this table will be a node in the network. These genes are filtered using a discoverable gene set. A network was built by calculating a matrix of Jaccard coefficients measuring the overlapping genes between these discoverable (or filtered) nodes. If there is a significant (according to user-supplied thresholds) overlap between the genes represented by the nodes, then there is a link between the nodes. Next, the workflow selects nodes with a keyterm in their study title or trait name and the nodes’ 1st nearest neighbors under the assumption that neighbors with significant overlap are also of interest. All other nodes are removed from the network, and the list of genes of the filtered network model serves as the trait-related gene set.

### Module-4: Over-representation analysis for genetic predisposition

Module 4 calculates the p-value of an overrepresentation test on the intersection of the metabolite interacting genes from module-2 and genes of the trait/disease-related GWAS network from module-3. Over-representation is calculated using the hypergeometric test. The total population is all genes in the discoverable gene set. The number of successes is (disease) genes from the trait/disease-related GWAS network. The draw size is the number of metabolite interacting genes; observed successes are the GWAS genes within the draw. As we are conducting a single test, no correction method is needed. Significant over-representation suggests a putative causal relationship with the polymorphism containing gene loci and metabolite change, potentially predisposing to disease development.

## RESULTS

### metGWAS 1.0 platform case assessments

We applied the metGWAS 1.0 platform to two case studies to evaluate performance on real data. Case 1 and 2 are taken from published metabolomics-GWAS studies with paired metabolite and GWAS data. We treated the metabolite data of these metabolomics-GWAS studies as a standalone dataset and linked the metabolites to interacting genes from the HMDB. We also created trait/disease-related GWAS networks for the phenotypes of interest (using Jaccard coefficient ≥ 0.5 and FDR-corrected p-value ≤ 0.05 as threshold). We compared our findings to those of the study to determine if previously identified gene loci could be rediscovered. We also reported any novel metabolite to phenotype gene interactions. Novel gene associations between metabolites and disease are possible since the GWAS catalogue study populations are independent of the case study populations and assess a similar phenotype or trait (e.g., heart disease).

#### Case study 1

A metabolite dataset (provided in **supplementary material 2)** for case study 1 was extracted from supplementary table 1 of a metabolomics-GWAS of human plasma, designed to identify the link between the lipidome and cardiovascular disease (CVD) [17]. A total of 42 unique metabolites were identified with significant genetic associations (p-value of < 1.5×10^−9^)[17]. We excluded 3 of the 42 metabolites as they were not mappable to an HMDB ID (i.e., non-specific metabolites belonging to total ceramides, total sphingomyelins, and total phosphatidylethanolamines). For the 39 remaining metabolites, the study reported associations with 11 genes, 5 of which were also associated with CVD.

We applied metGWAS to the 39 metabolites and identified the KEGG pathway glycerophospholipid metabolism as significantly enriched via MetaboAnalyst (**Table-1**). Next (in module 2), metGWAS mapped 86 interacting protein-coding genes to the glycerophospholipid metabolism pathway. This gene set serves as metabolite-interacting genes. Next (in module-3), metGWAS filtering of the full GWAS network with the key term “Cardiovascular disease” identified 84 nodes containing 189 genes. The 189 genes annotated to the cardiovascular disease-related GWAS subnetwork were then compared to the metabolite gene set by a hypergeometric test (in module 4). There is a significant overlap of metabolite-interacting human genes of the “glycerophospholipid metabolism” pathway with the cardiovascular disease-related GWAS subnetwork (*p* = 0.0028). Specifically, the nine overlapping genes were APOA5, PLA2G5, PLA2G2D, PLA2G2E, PLA2G2F, LRAT, PLA2G2A, PLB1, and PLA2G7. APOA5 was one of the genes identified in the original analysis [17] associated with CVD.

**Table-1:** The results of case study-1 using the metGWAS 1.0 platform.

| KEGG pathways for significant metabolites<br>(annotated metabolites, interacting genes) | Hypergeometric test | GWAS keyword |
| --- | --- | --- |
|  |  | Cardiovascular disease<br>(35 primary nodes, 49<br>neighbouring nodes; 189<br>genes) |
| Glycerophospholipid metabolism pathway<br>(14 HMDB ids, 86 genes) | # Overlapping genes | 9 |
|  | Overlapping gene names | **APOA5**, PLA2G5, PLA2G2D,<br>PLA2G2E, PLA2G2F, LRAT,<br>PLA2G2A, PLB1, PLA2G7 |
|  | P-value | 0.002838477 |

#### Case study 2

For case study 2, a metabolite dataset (provided in **supplementary material 2)** was extracted from supplementary table 2 of a publication on the relationship of urine metabolome with GWAS to investigate potential detoxification variants [18]. Case study 2 identified 56 unique metabolites with significant genetic associations (p < 2.9×10^−5^) [18] representing 5 genes.

Using MetaboAnalyst, the 56 metabolites enriched the four KEGG pathways (FDR≤ 0.05; **Table-2)** ‘aminoacyl-tRNA biosynthesis’, ‘glyoxylate and dicarboxylate metabolism’, ‘glycine, serine and threonine metabolism, and ‘alanine, aspartate and glutamate metabolism. These represented between 5-10 metabolites mapping to 90-206 protein interactions. Interestingly, the gene AGXT2 was present in all four enriched pathways and was one of the five genes reported by the study with significant metabolite associations.

**Table-2:**
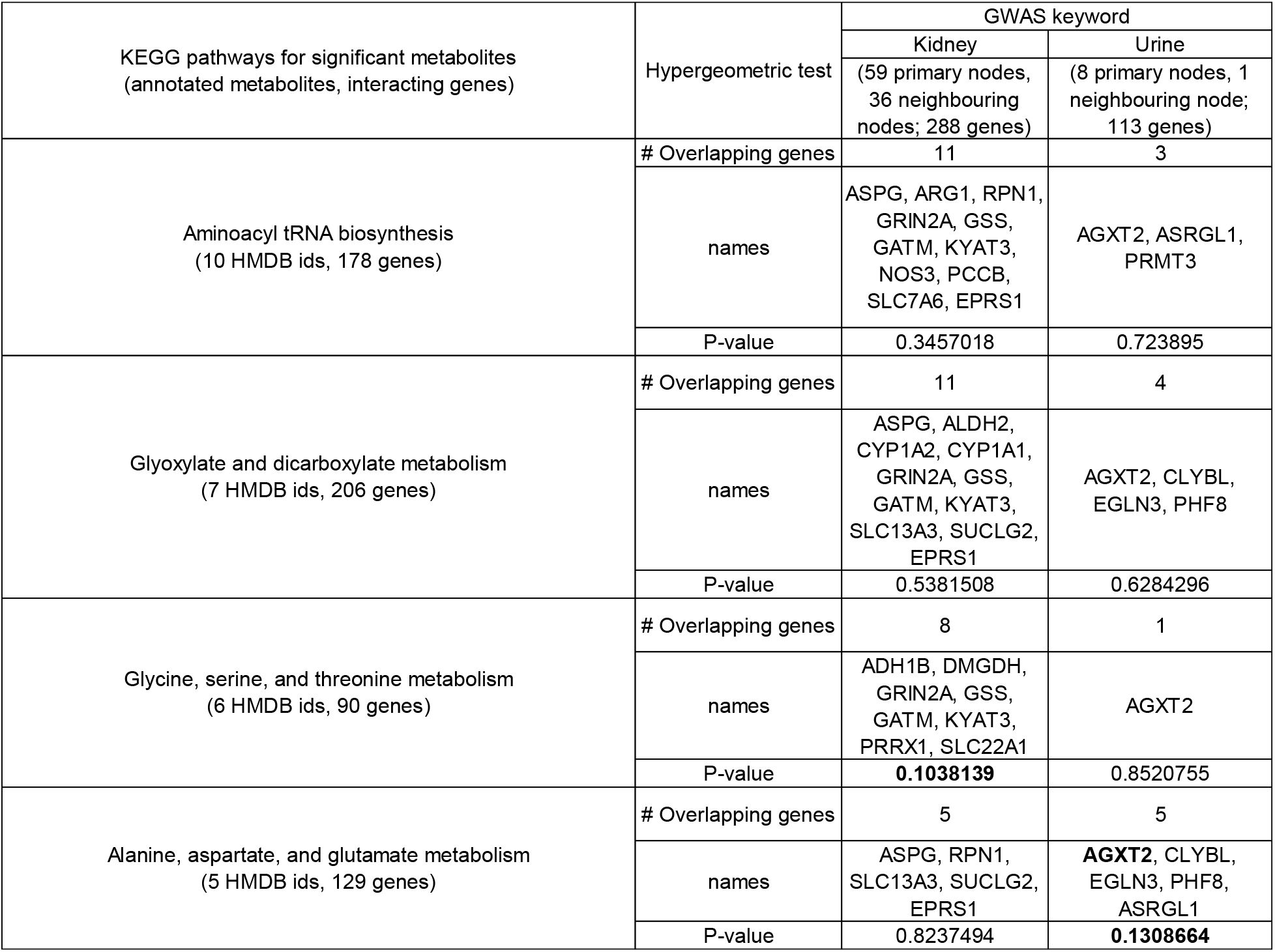
The results of case study-2 using the metGWAS 1.0 platform.

We selected the GWAS search keywords ‘Kidney’ or ‘Urine’ as they aligned with the metabolite study design of assessing urine to evaluate kidney detoxification capability. Search results identified a kidney-centred GWAS sub-networks of 95 nodes containing 288 genes (**Table 2**). The urine-centred subnetwork contained nine nodes encompassing 113 genes (**Table 2**). Eight genes overlap between the metabolite-interacting human genes of the “glycine, serine, and threonine metabolism” pathway and the kidney-specific GWAS network. A hypergeometric test found this to be marginal (*P* = 0.103). The overlapped genes are ADH1B, DMGDH, GRIN2A, GSS, GATM, KYAT3, PRRX1, and SLC22A1.

Assessing the metabolite-interacting human genes of the “alanine, aspartate, and glutamate metabolism” pathway found five genes (AGXT2, CLYBL, EGLN3, PHF8, and ASRGL1) overlapping to the urine-specific GWAS network. However, the hypergeometric test was marginal (*P* = 0.13) (**Table 2**). Interestingly, among the overlapping genes was AGXT2, also identified in the original case study as significantly associated with variation of metabolites in urine.

#### Primary and First Neighbor Nodes

To understand how the first-neighbour approach facilitated the metGWAS method, we looked at how primary and first neighbour nodes contributed to the intersecting metabolic and GWAS gene sets. In case study 1, all genes were derived from 1^st^ neighbour nodes, indicating that GWAS studies annotated as a cardiovascular disease were not the source of metabolic genome interactions. In contrast, the intersecting genes of case study 2 were all derived from the primary nodes. This suggests no new information was obtained by including the first neighbour nodes.

## DISCUSSION

A metabolomics-GWAS study is designed to identify the interaction of polymorphic variants with expression levels of a metabolite using paired samples. A series of logistic regression models identify gene variants influencing metabolite levels. Many metabolomics-GWAS studies do not use a case-control cohort design but instead utilize a population of healthy individuals to observe the natural variation. It is proposed that normal variants related to changes in metabolite levels can compound with other factors (e.g., other gene variants, environmental factors, etc.) and reach a tipping point leading to disease. The advantage of an integrated metabolomics-GWAS is that both direct and indirect associations may be observed.

Our metGWAS 1.0 platform uses the extensive resource of independent GWAS and metabolomics studies focused on similar diseases. As these samples are unpaired, we cannot utilize the same procedure as a standard metabolomics-GWAS. Instead, we focus on annotated protein-metabolite interactions as a directly interpretable finding. Our approach should identify generalizable results as both the metabolite and GWAS study population are independent.

In the first case study, metGWAS successfully reidentified one gene and identified other potentially novel metabolites to gene associations. The second case study identified associations with a marginal p-value near 0.1. The case studies presented essential differences in study design. Study 1 investigated a pathology where the phenotype to gene associations are likely stronger. In study 2, all participants were considered normal, and the variation in urine metabolome does not indicate pathology. The GWAS catalogue-derived network mainly comprises disease or phenotypes, while case study 2 did not contain any specific phenotype (i.e., healthy people). The polymorphisms found in natural variation may not be significantly associated with any pathology state. Urine is more challenging for the direct association as secreted metabolites may not be primarily produced by the kidney, with the liver being a major detoxification organ. While in case study 1, the assessment of CVD phenotype associations is more likely to produce overlapping results with the GWAS network that contains similar study designs. This was our observation with a significant overlap (p<0.05) that included a gene from the original study (APO5A).

Despite the marginal result for case study 2, some interesting findings are present. Of the five gene loci identified (i.e., AGXT2, CLYBL, EGLN3, PHF8, and ASRGL1) of the “alanine, aspartate, and glutamate metabolism” pathway, AGXT2 (i.e., reidentified gene from the original study) may be of interest. Within the GWAS Catalog, 16 GWAS studies identified 22 associations of AGXT2 with chronic kidney diseases using urinary metabolite measurements. Therefore, the AGXT2 gene loci may be an important candidate for further analyses of human detoxification capacity. The metGWAS platform may help narrow down the possible genetic loci for further study.

A critique of GWAS and metabolomics-GWAS is low reproducibility between different genetic populations. For example, Suhre and colleagues (case study 2) identified five gene loci as significant in a German population (i.e., SHIP study and KORA study) **[18]**. Schlosser and colleagues replicated four metabolite-gene variant associations using another German cohort (i.e., GCKD study) **[19]**. However, Flitman and colleagues could not identify any of Shure’s identified gene loci in a cohort from Lausanne, Switzerland **[20]**. It is equally challenging for the metGWAS 1.0 platform since it always executes its analyses using independent GWAS and metabolite cohort populations. Moreover, the GWAS database includes different population-based studies. Therefore, the reidentification of the same gene loci of a particular metabolomics-GWAS study using the metGWAS 1.0 platform is unlikely. However, we have shown here both case studies identified a gene loci from the original metabolomics-GWAS study.

Population variation is a well-known factor that undermines the reproducibility of GWAS. This may indicate why we had limited success in rediscovering the original metabolite to gene associations. This may also be advantageous as different SNPs and genes may drive the same metabolic shifts between study populations. This is supported by case study 1, where many statistically significant novel metabolites to SNP associations were identified.

Metabolite-GWAS studies directly measure SNP associations with metabolite levels on paired data. However, our metGWAS platform functions at the gene level. This may challenge polymorphism to gene mapping of distal non-coding polymorphisms. GWAS studies often yield SNPs in non-coding regions [21], which are mapped to the nearest gene, but SNPs can also have long-distance regulatory effects [22]. Additionally, metGWAS focuses on direct protein interactions with metabolites, meaning that protein-coding and non-coding genes that function upstream or downstream from the metabolite of interest will not be identified even though they may affect metabolite levels.

Using first neighbour nodes may bring irrelevant studies as the study phenotypes or traits do not contain the keyword search terms. However, the neighbours do have a significant gene overlap, suggesting some relationships. Assessing the first neighbours may be a tool to guide a literature search and determine previously unknown genetic relationships between phenotypes. Curiously, we found that the cardiovascular disease study results were only derived from 1^st^ neighbour node, while the urine metabolism study results were only from primary nodes. A likely reason for these opposite findings may lie in the selected GWAS subnetwork study designs and the metabolic studies. In case study 1, the primary cardiovascular disease nodes were connected to 1^st^ neighbour nodes representing cholesterol, blood glucose, or insulin dysregulation traits. These measurements are all related to metabolic syndrome, which can increase CVD risk [23]. In case study 2, we observed only the primary nodes contributing to both the urine and kidney anchored GWAS subnetworks. This suggests the primary nodes are more related to the metabolic genetic origin, and those first neighbour nodes may relate to non-metabolic genetic contributions.

## CONCLUSION

The metGWAS 1.0 platform showed success as a discovery platform for integrating metabolite and GWAS datasets. Our results indicate that associating studies related to specific pathologies was more successful than assessing normal variation. The network view of GWAS may lead to finding novel genetic associations between seemly unrelated phenotypes.

## AVAILABILITY AND REQUIREMENTS

**Project name:** metGWAS 1.0

**Project home page:** N/A

**Operating system(s):** Windows/Unix/Linux

**Programming language:** R

**Other requirements:** Details in the tutorial document, found in supplementary material 1

**License:** Free

**Any restrictions to use by non-academics:** N/A

## DATA AND CODE AVAILABILITY

Codes are available in supplementary material 1.

## SUPPLEMENTAL INFORMATION

***Supplementary material 1:*** An R-script with the user manual, split into two documents: a tutorial and a technical, for running the workflow, has been attached in this supplemental section. It also contains the necessary document and R-script for set-up the environment (i.e., q value and Bioconductor install.R, and RSelenium Driver SetUp.R) and running the workflow (i.e., objects folder).

***Supplementary material 2:*** The details of the case study-1 and case study-2 analysis and their results are provided here.

## ACKNOWLEDGMENTS

These studies are supported by research grants from the Canadian Institutes of Health Research (CIHR), FRN 143219 and Diabetes Canada (MBW). SRK was supported by a Diabetes Canada post-doctoral fellowship followed by a Banting and Best Diabetes Centre post-doctoral fellowship. BJC is supported by a Tier II Canada Research Chair. AO is supported by a CGS-M NSERC scholarship. The funders had no role in study design, data collection, and interpretation, or the decision to submit the work for publication.

## AUTHOR CONTRIBUTIONS

SRK, BJC, and MBW conceptualized and designed the research work executed by SRK and AO. SRK wrote the manuscript, edited by AO, EPG, BJC, and MBW. All authors reviewed the manuscript and gave final approval of the version to be published.

